# Short communication: automatic real-time monitoring of fungal spores: the case of *Alternaria* spp

**DOI:** 10.1101/2022.08.03.500168

**Authors:** Sophie Erb, Alexis Berne, Nina Burgdorfer, Bernard Clot, Marie-José Graber, Gian Lieberherr, Christine Sallin, Fiona Tummon, Benoît Crouzy

## Abstract

We present the first implementation of the monitoring of airborne fungal spores in real-time using digital holography. To obtain observations of *Alternaria* spp. spores representative of their airborne stage, we collected events measured in the air during crop harvesting in a contaminated potato field, using a Swisens Poleno device. The classification algorithm used by MeteoSwiss for operational pollen monitoring was extended by training the system using this additional dataset. The quality of the retrieved concentrations is evaluated by comparison with parallel measurements made with a manual Hirst-type trap. Correlations between the two measurements are high, especially over the main dispersion period of *Alternaria* spp., demonstrating the potential for automatic real-time monitoring of fungal spores.

---

The automatisation of operational pollen monitoring networks has been taking place over the past few years [1–3]. Recent instruments deliver airborne pollen concentrations in real-time at high-temporal resolutions, better corresponding to the active dynamics of meteorology that influences particle dispersion. This is resulting in a paradigm change in terms of development of modelling frameworks and communication to the public. In particular, this enables the integration of measurements directly into numerical weather models, which in turn deliver forecasts of airborne pollen concentrations at a spatial resolution on the order of one kilometre. Traditional monitoring networks have been based on the Hirst method [4]: microscope counting of particles collected on a band by impaction [5]. In Switzerland and many other regions, limited human resources do not allow for the routine counting of fungal spores. However, the impact of fungal spores on human health [6–8] and agriculture [9–11] is massive. Delivering accurate information on airborne spore concentrations would therefore allow improved prevention and treatment of respiratory allergies and a more targeted use of fungicides in agriculture, which in turn would reduce costs and environmental impacts.

In 2019, the Swiss Federal Office of Meteorology and Climatology MeteoSwiss started deploying the first national automatic pollen monitoring network based on the Swisens Poleno [12]. Operational particle identification and counting is performed using reconstructed images taken in flight using digital holography (see Figure 1 and [13]). In the Swisens Poleno, each particle sufficiently large (i.e. larger than *∼* 5*µm* since one pixel on the image corresponds to 0.595 *µm* in the physical domain) to be resolved by digital holography is imaged by two orthogonal cameras. A two-stage classifier is then used to identify each particle. In the first stage, pollen is distinguished from other particles (e.g. dust, fibers, biological debris). In the second stage, a convolutional neural network is then applied to discriminate between different allergenic pollen taxa. Extending the scope of the device to the identification of other types of (bio-)aerosol is feasible under two conditions: first, the particles of interest need to fall within a suitable size range and second, correctly labelled reference images must be available for training the classifier.

**Fig 1.**
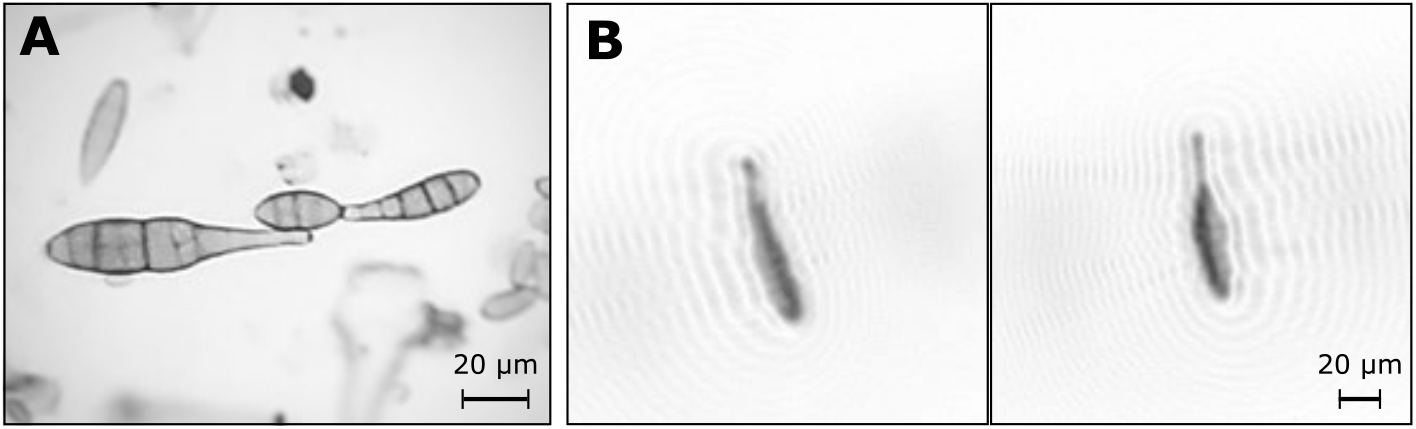
A. Microscopic image of *Alternaria* spores (600x magnification), B. Holographic images of a single *Alternaria* spore taken in flight with the Swisens Poleno.

Pathogens of the genus *Alternaria* are responsible for several crop diseases, commonly referred to as early blight, which affect a wide range of crops (e.g. potatoes, tomatoes, carrots). *Alternaria* species present various degrees of specialisation regarding their host plants. Yet, they share common morphological traits suggesting a potential for genus-level identification with operational monitoring devices. Specifically, as far as the size is concerned (20 - 200 *µm* [14]), *Alternaria* spp. spores are large enough to be imaged by the Swisens Poleno (see Figure 1).

Extending the potential of the classifier [12] to the genus *Alternaria* requires a clean dataset of spores measured by the Swisens Poleno. To this end, we identified an event with extremely high *Alternaria* concentrations (see Figure 2.A, 26.07.2021) using routine Hirst-based measurements [5] taken at MeteoSwiss in Payerne, Switzerland. Using concurrent meteorological observations of wind direction and speed data, we were able to link this event with the harvest of a neighbouring potato field infested with *Alternaria solani*. Since the collection of airborne particles occurs on a clock-controlled rotating drum, we can associate the position of the particle on the band to the time at which it impacted (Figure 2.A). Considering that the Swisens Poleno devices are collocated with the Hirst measurements (as described in [15]), the peaks in the total number of particles measured by the Swisens Poleno could be matched with the Hirst measurements of *Alternaria* spores (see Figure 2.B). We isolated the corresponding peak events in the Swisens Poleno data, cleaned them by manually selecting the events presenting correct holographic images and obtained a dataset of 2767 events with a morphology (size and shape) compatible with *Alternaria*. Figure 1.B shows an example of such an event (consisting of two holographic images taken at 90° from one another). We added a class for *Alternaria* and straightforwardly retrained the neural network originally developed by [12].

**Fig 2.**
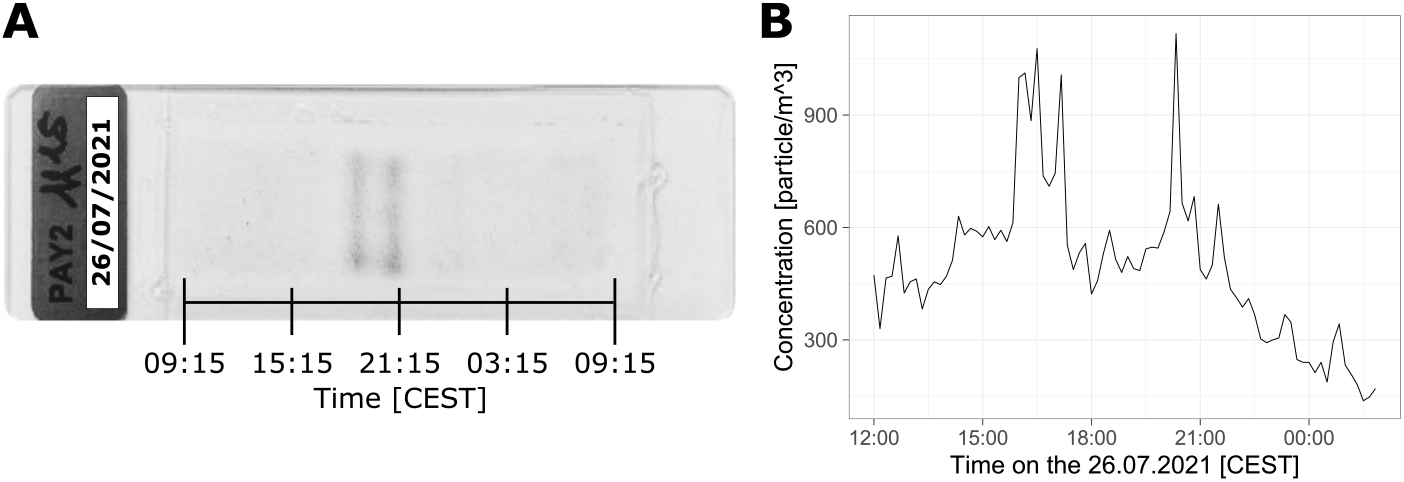
A. Hirst slide with peak particle concentrations appearing as dark vertical lines, B. The corresponding total number of particles measured by the Swisens Poleno, run in parallel to the Hirst impactor on the roof of the MeteoSwiss station, Payerne, Switzerland.

To validate the newly updated classifier, we ran it on a period from 3 March to 21 September 2021. This was made possible by the fact that raw data produced by the Swisens Poleno (reconstructed images) is systematically archived at MeteoSwiss. As described in [12], we used thresholds on the outputs of the neural network to limit false-positive detection. In the absence of established guidelines for fungal spores, we used a value of 0.8 as threshold since this provides good results for operational pollen monitoring. Absolute scaling with the automatic system was obtained using the procedure described in [1]. The scaling factor was computed over the dispersion period of *Alternaria* (Figure 3.B) to limit the effect of false-positive detection. The scaling factor used for the automatic raw particle count is 0.115, resulting from the higher sampling of the Swisens Poleno compared to the Hirst impactor. Manual counting of *Alternaria* spores is labour intensive and round-the-year counting of spores is not feasible. We therefore selected three days per month (21 days in total) representing various *Alternaria* concentrations for which manual counts were made. These were compared with the concentrations from the newly-trained classifier (Figure 3.A). Nevertheless, an arbitrary choice of 21 days for comparison hardly qualifies as a comprehensive validation. We therefore performed additional continuous counts for a period of 30 days (Figure 3.B). These results show that for the days with higher concentrations, the Swisens Poleno tends to underestimate concentrations, possibly suggesting that saturation of the device occurs. Hence, we checked the total particle counts of the Poleno on days with high peaks of particles and removed two days (27 July and 12 August) which presented saturation to compute correlations and the scaling factor for comparing the two time series’. In more recent versions of the instrument this issue has been corrected by the introduction of adaptive sampling rates. To quantify the correlation between the manual and automatic time series’, we used Spearman’s rank correlation due to the non-normality of the data and the increased robustness of this metric to outliers. As a complement, Pearson’s correlation (disregarding the two saturated days) is also computed. Altogether, considering the differences between manual and automatic systems, from the sampling (impaction vs. Sigma 2 inlet) to the identification method (optical microscope counting vs. digital holography), we observe a solid agreement between the two methods. Notice that the Spearman value is lower (*ρ* = 0.76) over the extended period (Figure 3.A) than over the continuous period (*ρ* = 0.91) (Figure 3.B), suggesting the potential for improvement of avoiding false-positives. Finally, even though we trained the model on a dataset specific to a single *Alternaria* species, we obtained reasonable results for the whole genus. Nonetheless, we expect to improve *Alternaria* recognition using a more general training dataset extended to other *Alternaria* species.

**Fig 3.**
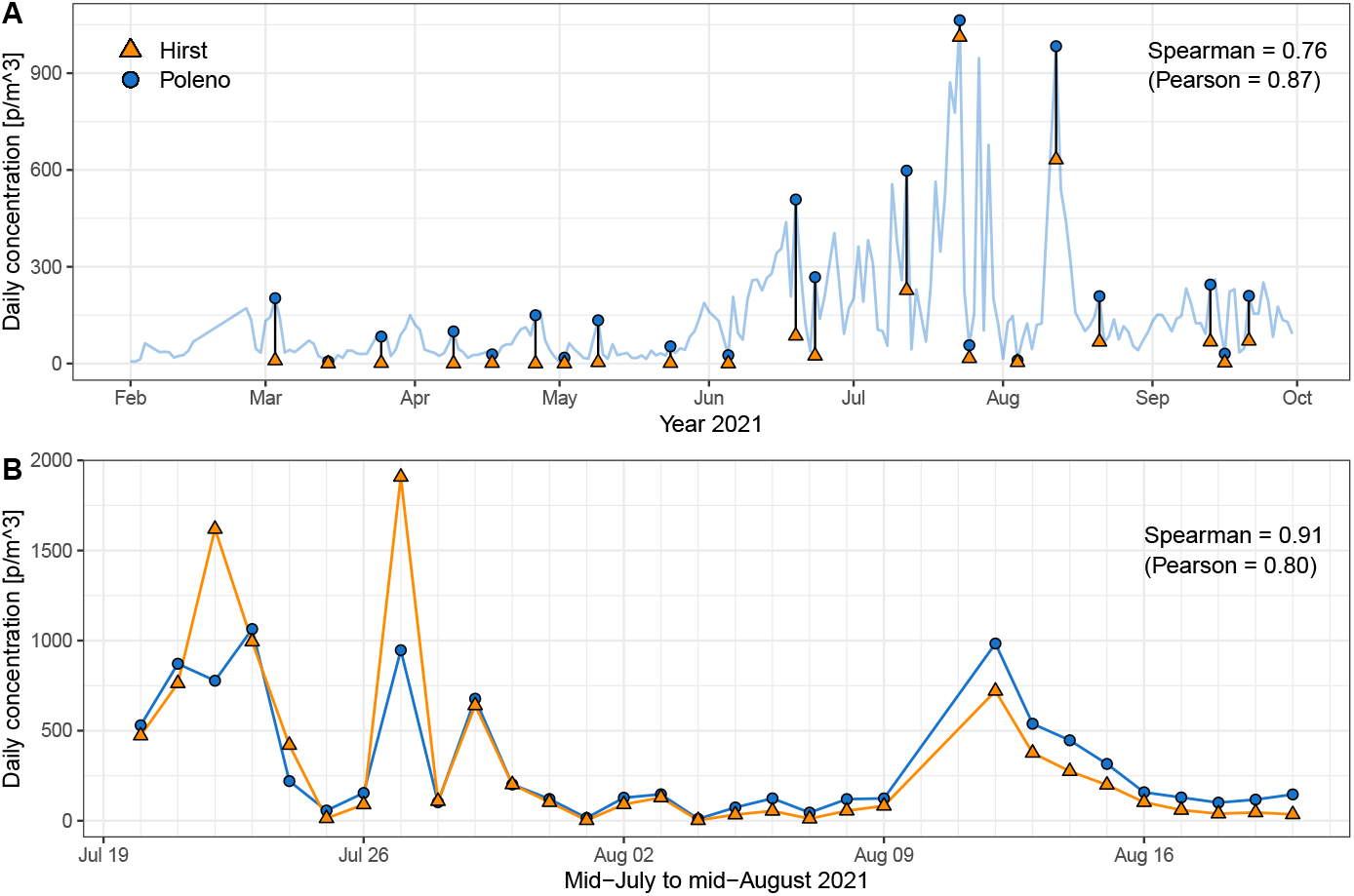
A. Comparison between manual (Hirst) and automatic (Swisens Poleno) pollen measurements for 21 days sampled from March to September 2021. Dates were chosen a priori based on the concentration time series of the Swisens Poleno only (light blue line). Three days per month were sampled, each representing the lowest and highest values as well as a medium value. B. Comparison between manual and automatic fungal spores concentration measurements for a month between 20 July and 20 August 2021.

This first comparison paves the way for the monitoring of other fungal spores with the aim of supporting the future development of adapted agricultural methods with targeted use of fungicides. Furthermore, the availability of such operational monitoring data at high temporal (hourly) resolution will serve to improve our understanding of the role of fungal spores in triggering allergic reactions and to improve both diagnosis and treatment thereof. Moreover, it will allow the determination of critical exposure thresholds for the allergic response of sensitised individuals.

## Authors’ contributions

SE conducted the study, NB contributed to the neural network architecture, MJG and CS provided the manual Hirst counts, AB, BClot, GL and FT contributed to writing and BCrouzy supervised the study and contributed significantly to writing.

